# Deciphering the Immune Subtypes and Signature Genes: A Novel Approach Towards Diagnosing and Prognosticating Severe Asthma through Interpretable Machine Learning

**DOI:** 10.1101/2024.04.15.589644

**Authors:** Yue Hu, Yating Lin, Bo Peng, Chunyan Xiang, Wei Tang

## Abstract

Asthma, a pervasive pulmonary disorder, affects countless individuals globally. Characterized by chronic inflammation of the bronchial passages, its symptoms include cough, wheezing, dyspnea, and chest tightness. While many manage their symptoms through pharmaceutical interventions and self-care, a significant subset grapples with severe asthma, posing therapeutic challenges. This study delves into the intricate etiology of asthma, emphasizing the pivotal roles of immune cells such as T cells, eosinophils, and mast cells in its pathogenesis. The recent emergence of monoclonal antibodies, including Mepolizumab, Reslizumab, and Benralizumab, offers therapeutic promise, yet their efficacy varies due to the heterogeneous nature of asthma. Recognizing the potential of personalized medicine, this research underscores the need for a comprehensive understanding of asthma’s immunological diversity. We employ ssGSEA and LASSO algorithms to identify differentially expressed immune cells and utilize machine learning techniques, including XGBoost and Random Forest, to predict severe asthma outcomes and identify key genes associated with immune cells. Using a murine asthma model and an online database, we aim to elucidate distinct immune-centric asthma subtypes. This study seeks to provide novel insights into the diagnosis and classification of severe asthma through a transcriptomic lens.

## Introduction

Asthma, a ubiquitous pulmonary malaise, affects millions of people worldwide^1^. As an enduring inflammatory perturbation of the bronchial passageways, it exhibits symptoms encompassing chronic cough, wheezing, dyspnea, and thoracic tightness^2^. These clinical presentations originate from the inflammatory cascade within the airways, precipitating phenomena such as mucosal overproduction, architectural transformation of the airway walls, and amplified bronchial susceptibility^3^. While a predominant fraction of individuals with asthma can adeptly regulate their clinical manifestations through judicious pharmaceutical strategies and astute self-management, there exists a significant cohort persistently besieged by the severe manifestation of this condition, presenting a vexing therapeutic enigma^4–6^. Severe asthma is delineated as a malady demanding augmented doses of inhaled corticosteroids concomitant with ancillary controller medicaments or systemic corticosteroids to thwart unbridled exacerbations, or one that remains obdurate despite such pharmacological expeditions^7^. The complex therapeutic landscape of severe asthma, marked by patient diversity, places a significant burden on those affected and presents a challenging puzzle for healthcare providers to solve^8^. Identifying key diagnostic markers and understanding the subtle pathophysiological aspects of severe asthma is crucial for developing personalized treatments.

Asthma’s etiology is complex, comprising a network of immune cells and signal pathways. T cells are crucial, anchoring the immune system’s coordination^9^. In asthma, these cells become active, producing a slew of cytokines that alter immune communication^10^. Specifically, T helper 2 cells leave a lasting impact by releasing key cytokines—interleukin-4, −5, and −13—which drive inflammation and energize the immune front lines^11,12^. Eosinophils, crucial in fighting parasites, swell in number, exacerbating inflammation through cytokines and other agents, attacking the delicate airway lining and summoning more immune cells to the site^13,14,15^. Mast cells, central to allergic reactions, contribute to the development of asthma by releasing inflammatory agents like histamine, heightening airway disruption and sensitivity^16,17^. In essence, asthma manifests as a complex immunological puzzle with diverse elements like T cells, eosinophils, and mast cells. Each plays a distinct role in the cascade of inflammation and sensitivity. Unraveling the complex molecular patterns and signaling pathways in asthma is critical for developing new treatments to address this persistent condition.

Recent advancements in asthma treatment have been marked by the introduction of novel monoclonal antibodies, including Mepolizumab, Reslizumab, and Benralizumab. These agents specifically target immune pathways associated with IL-5 and have shown promising outcomes for select patient groups. For example, while early Mepolizumab trials were not fully satisfactory, later research in patients selected based on biomarkers revealed significant improvements in lung function and reduced reliance on corticosteroids^18^. Nonetheless, this optimism is not universally applicable due to the significant heterogeneity observed among asthma patients, especially in type 2-high asthma, evidenced by the mixed responses to anti-IgE treatments such as omalizumab, with about 70% of recipients showing improvement^19^.

In the thrust towards personalized medicine, recognizing the diverse immunological foundations of asthma is critical. The genomic revolution underscores the urgency to establish a comprehensive omics framework to identify reliable biomarkers for severe asthma. A detailed examination of immune-related signature genes and precise categorization of immune molecular subtypes are crucial steps towards providing individualized treatment strategies for asthma. The classification of asthma includes varied subtypes like allergic, non-allergic, occupational, aspirin-exacerbated respiratory disease, potentially fatal, exercise-induced, and cough variant asthma, each requiring specific clinical approaches. These subtypes display a wide array of clinical presentations and underlying mechanisms, posing challenges to patient management and ongoing care. The EAACI guidelines offer clear definitions for allergic and eosinophilic asthma, classifying them as distinct subtypes of T2-high diseases^20^. However, the sharp distinction between allergic and eosinophilic asthma may be inadequate due to significant overlap in patients, highlighting the need for more precise and scientific classification^21,22^.

The ssGSEA method, commonly used in immune cell infiltration analysis, estimates the relative enrichment of specific gene sets (immune cell gene sets) in samples based on gene expression data^23^. LASSO is a regression analysis method that performs variable selection and regularization to enhance prediction accuracy and model interpretability. Gene studies involving numerous variables often face reproducibility issues. LASSO helps in selecting genes consistently associated with outcome variables to build replicable models^24^. We aim to dissect the immune complexities in patients with severe asthma, utilizing ssGSEA and LASSO to identify differentially active immune cells. By focusing on immune-associated differential genes and applying machine learning techniques like XGBoost, Random Forest, and LASSO, we strive to predict severe asthma outcomes and identify key genes closely related to immune mechanisms. Validating this genetic mosaic, we’ve summoned the insights of an asthmatic murine model and perused an ancillary online compendium. By exploring the diverse functions, pathways, immune cell infiltrations, and immunological characteristics, we aspire to provide groundbreaking insights that will enhance the diagnosis and detailed categorization of severe asthma via transcriptomics.

## Materials and Methods

### Raw Data Acquisition and Preprocessing

Gene expression profiles for asthma research were obtained from four distinct datasets: U-BIOPRED, Arron JR et al., Li Q et al., and Baines KJ et al. ^25,26,27^, each with a unique GEO accession number. These datasets include RNA from healthy controls, moderate and severe asthma cases, with the sample sizes and analysis platforms varying across studies. Detailed information on the datasets, including sample counts and microarray platforms used, can be found in Table S3.

### Assessment of Immune Cell Infiltration

To analyze immune cell infiltration, we applied single-sample gene set enrichment analysis (ssGSEA) to determine the abundance of 28 immune cell subtypes in individual samples, using enrichment scores to quantify each subtype’s relative proportion. Statistical significance was assessed with the Wilcoxon rank sum test, considering p-values below 0.05.

For identifying critical immune cell subsets across patient groups, we utilized LASSO regression, dividing our data into training and validation sets. Optimal parameters were determined via 10-fold cross-validation, selecting the penalty parameter (λ) that minimized cross-validation error. Immune cell subsets with non-zero coefficients after LASSO regression were further analyzed.

### WGCNA Network Construction and Module Identification

We used the WGCNA package to create a co-expression network from gene expression data, applying soft-thresholding to achieve a scale-free topology^28^. After determining the ideal soft-thresholding power, we constructed a topological overlap matrix and performed hierarchical clustering to detect gene modules. Correlations between these modules and immune cell subtypes were calculated using ssGSEA scores, focusing on modules with the strongest correlations for further analysis.

### Evaluation of Immune-Associated Differential Genes

In this study, we procured 2483 immune-related genes from the Immunology Database and Analysis Portal (ImmPort) and retrieved 1378 immune genes from the InnateDB database. To identify differentially expressed genes (DEGs) between control and severe asthma samples, we employed the “limma” R package with criteria of | log2 (fold change) | > 0.5 and a false discovery rate (FDR) of 0.05. We ascertained immune-associated DEGs by intersecting immune-related genes from ImmPort and InnateDB, DEGs, and genes in feature modules.

### Enrichment of Biological Functions and Pathways

We utilized the R package “clusterProfiler”(version 4.4.4) to conduct Gene Ontology (GO) biological functions and Kyoto Encyclopedia of Genes and Genomes (KEGG) pathway enrichment analyses^29^. GO enrichment analysis identified biological functions enriched among input genes, while KEGG pathway enrichment analysis revealed associated enriched pathways. Significantly enriched biological functions and pathways were chosen based on a threshold of p-value less than 0.05 and an FDR less than 0.05. Additionally, we generated a visual representation of enriched biological functions and pathways using the R packages ggplot2 (version 3.3.6) and GOplot (version 1.0.2).

### Machine Learning Models for Sophisticated Feature Selection and Visualization

We divided the combined dataset, comprising 114 samples (21 healthy controls and 93 severe asthma cases), evenly into training and test sets using the caret package in R. We then employed Lasso regression, XGBoost, and Random Forest algorithms for model building and feature selection. Lasso regression, known for its capacity to reduce the number of features, was used for its ability to enhance model interpretability. XGBoost, a gradient-boosting framework, was chosen for its performance in classification tasks, and Random Forest was selected for its robustness and effectiveness in handling complex interactions and classifications. The ROC curve and AUC were calculated to evaluate the models’ ability to distinguish between the two groups, demonstrating the practical utility of these algorithms in selecting and visualizing key features indicative of severe asthma.

### Unearthing Distinct Subgroups via Unsupervised Clustering

To identify subgroups within severe asthma patients, we used consensus clustering with the ConsensusClusterPlus package. The optimal number of clusters was determined using the consensus score matrix, CDF curve, PAC score, and Nbclust indicators. For visualization, we applied PCA to distinguish between the subtypes, with PCA plots highlighting the differences. Further details of this analysis are available in the Supplementary Information.

### Small-Molecule Compound Prognostication

We executed Connectivity Map (CMap) analysis to forecast small-molecule compounds targeting immune microenvironment subtypes 1 and 2, following previously established methods^30,31^. Succinctly, we procured 1309 drug signatures from the Connectivity Map database (CMap, https://clue.io/) and chose the top 150 upregulated and 150 downregulated expression profiles as input data. Employing the eXtreme Sum (XSum) algorithm, we computed CMap scores, and subsequently, the top five small-molecule compounds were designated for visualization.

### External Validation of Crucial Genes

We employed external datasets, comprising GSE74986, GSE137268, and GSE74075, to ascertain the capacity of immune related signature genes to differentiate severe asthma patients from healthy controls. Diagnostic effectiveness was visualized through AUC curves generated with the R package “pROC” (version 1.18.0).

### Mouse Model

The HDM-induced murine asthma paradigm was established as delineated previously^32^. Succinctly, WT mice were sensitized intranasally with 100 mg HDM in 40 mL PBS on Day 0 and confronted intranasally with 10 mg HDM from Days 7 to 11, culminating in harvest on Day 14.

### Real-time RT-PCR Analysis

Pulmonary tissue RNA was extracted employing AG RNA exPro RNA (AG21101). Total RNA underwent reverse transcription utilizing HiScript® III RT SuperMix for qPCR (Vazyme, Nanjing, China) for complementary DNA (cDNA) synthesis. The primers implemented for RT-PCR analysis were as Supplement Table S4.

The qRT-PCR was executed with ChamQ Universal SYBR qPCR Master Mix (Vazyme, Nanjing, China) and an Applied Biosystems QuantStudio 3 (Applied Biosystems, CA, USA). Relative quantification was ascertained against a standard curve, and specific values were normalized against murine GAPDH mRNA. The outcomes were characterized as a relative augmentation in mRNA expression in comparison to control values.

## Results

### 1. Differential Immune Cell Landscape in Severe Asthma Patients

Our study, outlined in Figure 1, systematically identified unique immune cell profiles and gene markers in severe asthma, culminating in the discovery of distinct patient subtypes and potential therapeutic targets through advanced multi-phase analysis and validation. In our initial exploration of the GSE76262 gene expression matrix, we conducted a Principal Component Analysis (PCA) across three categories: healthy control, moderate asthma, and severe asthma. Interestingly, the PCA revealed a marked proximity between the healthy control and moderate asthma groups, suggesting a high degree of similarity in their gene expression profiles. In stark contrast, the severe asthma group manifested a discernible separation from the other two, underlining its distinct gene expression landscape (Figure 2A).

**Figure 1.**
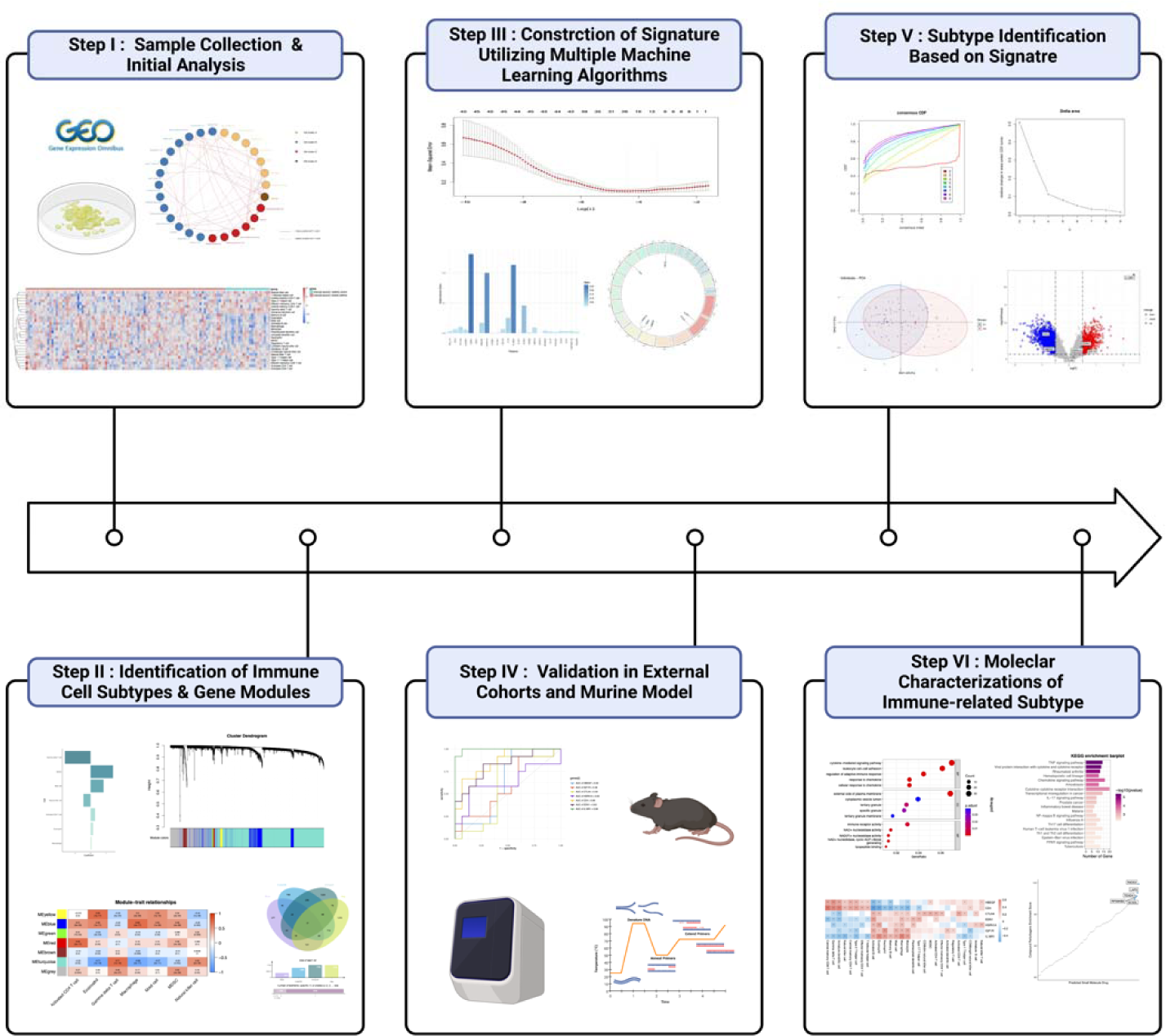
Flowchart Step I: Sample Collection & Initial Analysis. Collection of sputum samples from healthy controls and severe asthma patients, followed by immune cell abundance analysis using ssGSEA and PCA. This step sets the stage for identifying differences in immune cell profiles between the two groups. Step II: Identification of Immune Cell Subtypes & Gene Modules Utilization of LASSO regression analysis to identify key immune cell subtypes associated with asthma progression. Application of WGCNA to determine correlated gene modules, further refining the cellular and molecular landscape. Step III: Construction of Signature Utilizing Multiple Machine Learning Algorithms Implementation of machine learning algorithms like Lasso regression, XGBoost, and Random Forest to identify a robust set of key genes (e.g., HBEGF, HSPA1A, CD4, IL18R1, EDN1, CTLA4, and IGF1R) that can distinguish between healthy and severe asthma patients. Step IV: Validation in Outer Cohorts and Murine Model Validation of the identified key genes using ROC curves in test and validation sets, as well as external datasets. Further validation is conducted using a murine asthma model, confirming the genes’ roles in asthma pathogenesis. Step V: Subtype Identification Application of consensus clustering to identify distinct subtypes within severe asthma patients (C1 and C2). Exploration of differential gene expression and immune cell abundance between these subtypes to understand their unique characteristics. Step VI: Molecular Characterizations of Immune-related Subtype Detailed molecular characterization of the identified subtypes, focusing on differential gene expression and immune cell abundance. Recommendations for potential drug targets based on Cmap database analysis, paving the way for tailored therapeutic strategies.

**Figure 2:**
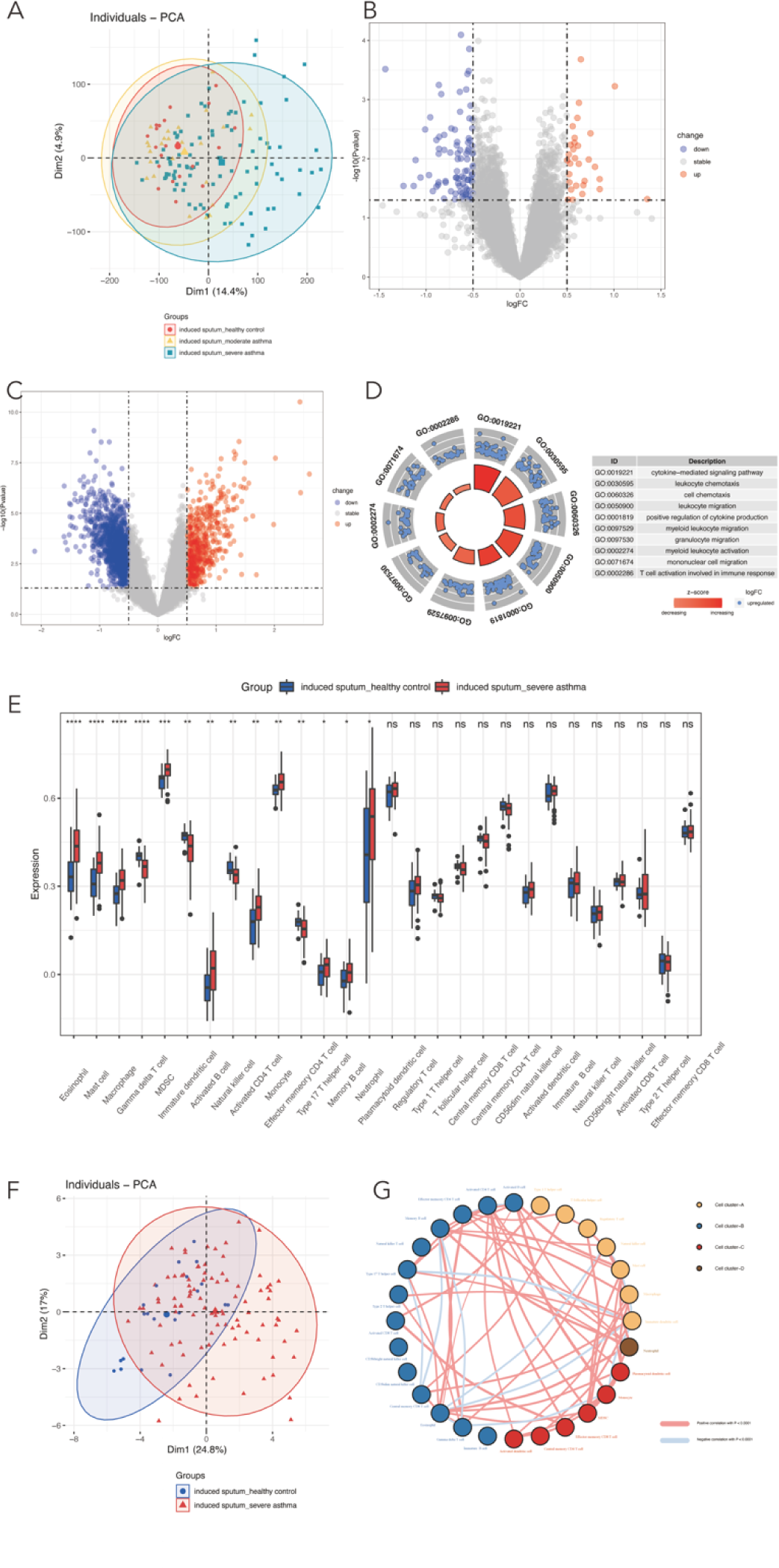
Immune cell abundance analysis in sputum samples from healthy controls and severe asthma patients. (A) PCA of gene expression from dataset GSE76262 delineates the distinction and overlap among healthy controls, moderate, and severe asthma. Notably, the proximity between healthy and moderate groups contrasts with the distinct separation of the severe asthma cluster. (B) Differential expression profiling between healthy controls and moderate asthma patients identifies 30 upregulated and 90 downregulated genes, underlining transcriptional similarities. (C) Extensive differential expression analysis between healthy controls and severe asthma patients reveals 593 upregulated and 1,493 downregulated genes, highlighting the pronounced molecular divergence in severe asthma. (D) Enrichment scores of immune cell types obtained using the ssGSEA method, highlighting the differences in immune cell abundance between healthy controls and severe asthma patients. (E) PCA plot of immune cell infiltration, illustrating substantial differences in immune cell abundance clustering between healthy controls and severe asthma patients. (F) Correlation heatmap of immune cell abundance coefficients, showcasing the relationships among immune cell types. (G) Heatmap visualization of the distinct immune cell composition in sputum samples from healthy controls and severe asthma patients.

Given the clinical challenges of managing severe asthma, we found it paramount to hone our research focus onto this group. A comparison between healthy controls and moderate asthma patients revealed only a small number of differentially expressed genes: 30 upregulated and 90 downregulated (Figure 2B). The limited gene differential suggests a closer transcriptional resemblance between these two groups as compared to the severe asthma cohort.

Considering the notable differences in gene expression patterns and the clinical implications of severe asthma, we proceeded with a more in-depth investigation into this category. Between the healthy control and severe asthma groups, our analysis discerned 593 upregulated and 1,493 downregulated genes (Figure 2C). This led to a GO biological process enrichment analysis, which highlighted significant immune-related activities, such as cytokine-mediated signaling and leukocyte migration, among the genes upregulated in severe asthma (Figure 2D). These findings underline the importance of immune cell migration and cytokine signaling in severe asthma. Given the pronounced enrichment of immune-related processes in severe asthma, it became paramount to delve deeper into this facet.

To further elucidate the immune contrasts between healthy individuals and severe asthma patients, we embarked on an investigation of the variations in enrichment scores across 28 immune cell subsets from the dataset using the ssGSEA method. Leveraging the ssGSEA method, we determined the abundance scores of immune cells present in sputum samples from both cohorts. Our inquiry revealed that severe asthma patients manifested increased enrichment scores for eosinophils, mast cells, macrophages, MDSCs, activated B cells, activated CD4 T cells (defined as CD4 T cells in an elevated state of responsiveness post-stimulation), monocytes, Th17 cells, memory B cells, and neutrophils when juxtaposed with healthy controls. Conversely, the γδT cells, immature DC cells, NK cells, and effector memory CD4 T cells exhibited reduced enrichment scores in the severe asthma samples (Figure 2E). It’s worth highlighting that while variations in some cell types, such as Th17 cells, may seem marginal, even these subtle distinctions could have profound implications in the intricate landscape of severe asthma pathogenesis^33^.

Another PCA focused on immune cell infiltration showed significant differences in the immune cell profiles of healthy versus severe asthma patients, further buttressing the premise that immune cell perturbations are deeply intertwined with the pathogenesis of severe asthma (Figure 2F). Further analysis of immune cell correlations revealed complex interaction patterns among the 28 immune cell types, identifying four unique interaction patterns (Figure 2G). This comprehensive view of immune cell interactions enriches our understanding of severe asthma’s pathogenesis.

### 2. Identification of Immune-Related DEGs

We employed LASSO regression analysis to pinpoint critical immune cell subtypes associated with severe asthma progression from the 28 discerned via ssGSEA (Figure 3A, 3B). Our findings spotlighted seven immune cell subtypes - MDSC, mast cell, activated CD4 T cell, eosinophil, macrophage, NK cell, and γδT cell, all of which had non-zero coefficients, underscoring their significance in asthma progression (Figure 3C). These cells play an integral role in asthma’s pathogenesis.

**Figure 3.**
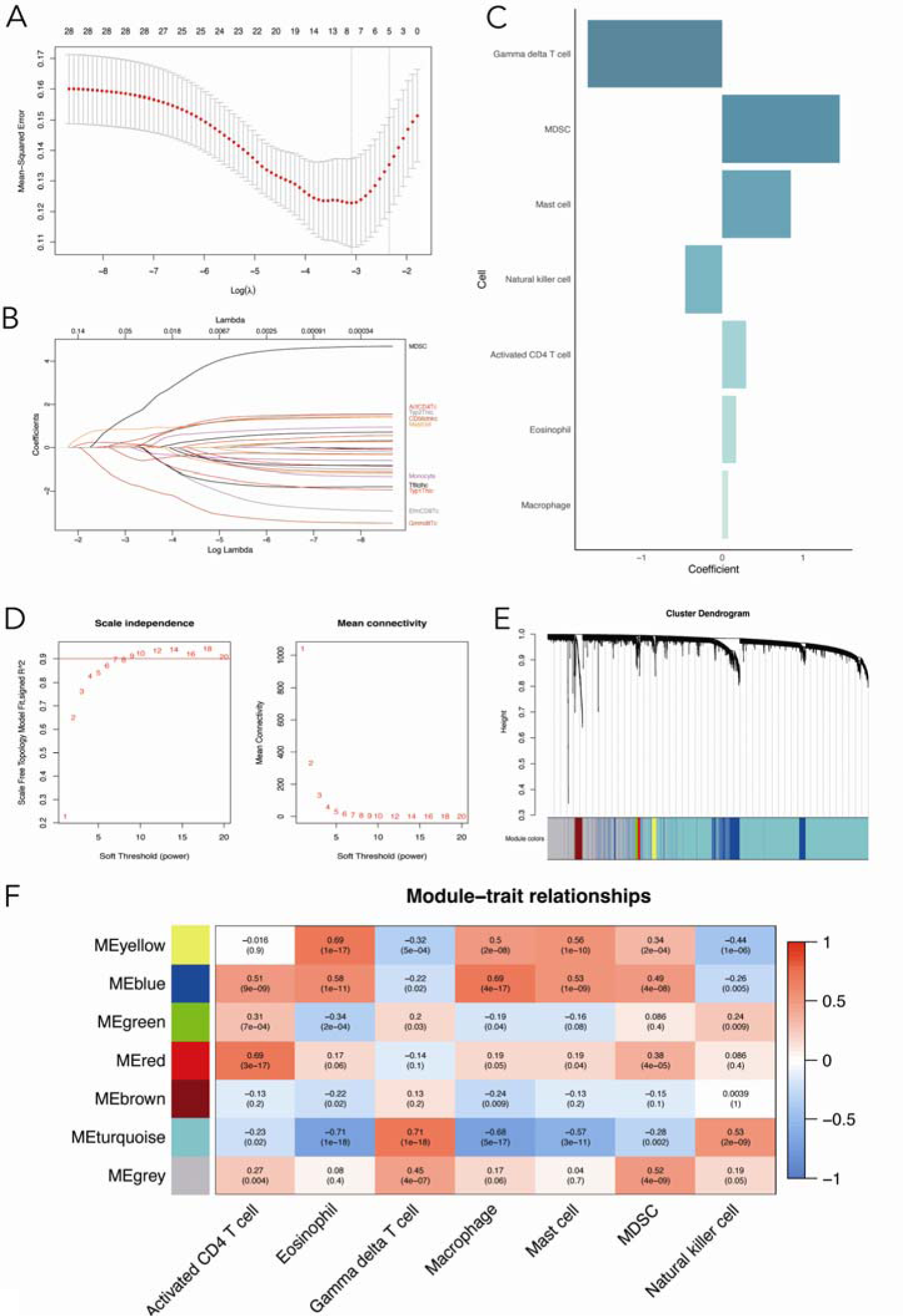
Identification of key immune cell subtypes and gene modules associated with asthma progression. (A-B) LASSO regression analysis to identify the most relevant immune cell subtypes for asthma progression based on the ssGSEA results. (C) The seven immune cell subtypes (MDSC, mast cell, activated CD4 T cell, eosinophil, macrophage, NK cell, and γδT cell) with non-zero coefficients, indicating their significant association with asthma progression. (D) Determination of the optimal soft-thresholding parameter for WGCNA using the PickSoftThreshold function. (E) Hierarchical clustering algorithm results, depicting the clustering tree divided into seven gene modules of different colors, each containing a set of highly correlated genes. (F) Heatmap showing the correlation between gene modules and the seven selected immune cell subtypes, with the Blue and Turquoise modules demonstrating associations with different sets of immune cell subtypes.

In a weighted gene co-expression network analysis (WGCNA) of sputum samples from both healthy controls and severe asthma patients, we identified core modules corresponding to these seven unique immune cell subtypes. The soft threshold β was determined at 14, establishing an optimal power value for coexpression network construction (Figure 3D). This resulted in seven gene modules, each marked by a distinct color and related genes, correlating with the immune cell subtypes (Figure 3E, 3F). Notably, the Blue module showed a positive correlation with several immune cells, whereas the Turquoise module exhibited a mix of negative and positive correlations with different cells, indicating their varied roles in severe asthma. Moreover, these genes were confirmed in authoritative immune databases as actively participating in immune regulation and response processes. This approach identified 28 immune-related DEGs from the Turquoise module and differentially expressed genes between healthy controls and severe asthma patients, both found in the ImmPort and InnateDB databases (Figure 4A). The Blue module yielded 17 such DEGs (Figure 4B). Key genes like IGF1R, CTLA4, HSPA1A, EDN1, and IL18R1 were upregulated in severe asthma, while HBEGF and CD4 were downregulated (Figures 4C, D), highlighting their critical role in asthma progression.

**Figure 4.**
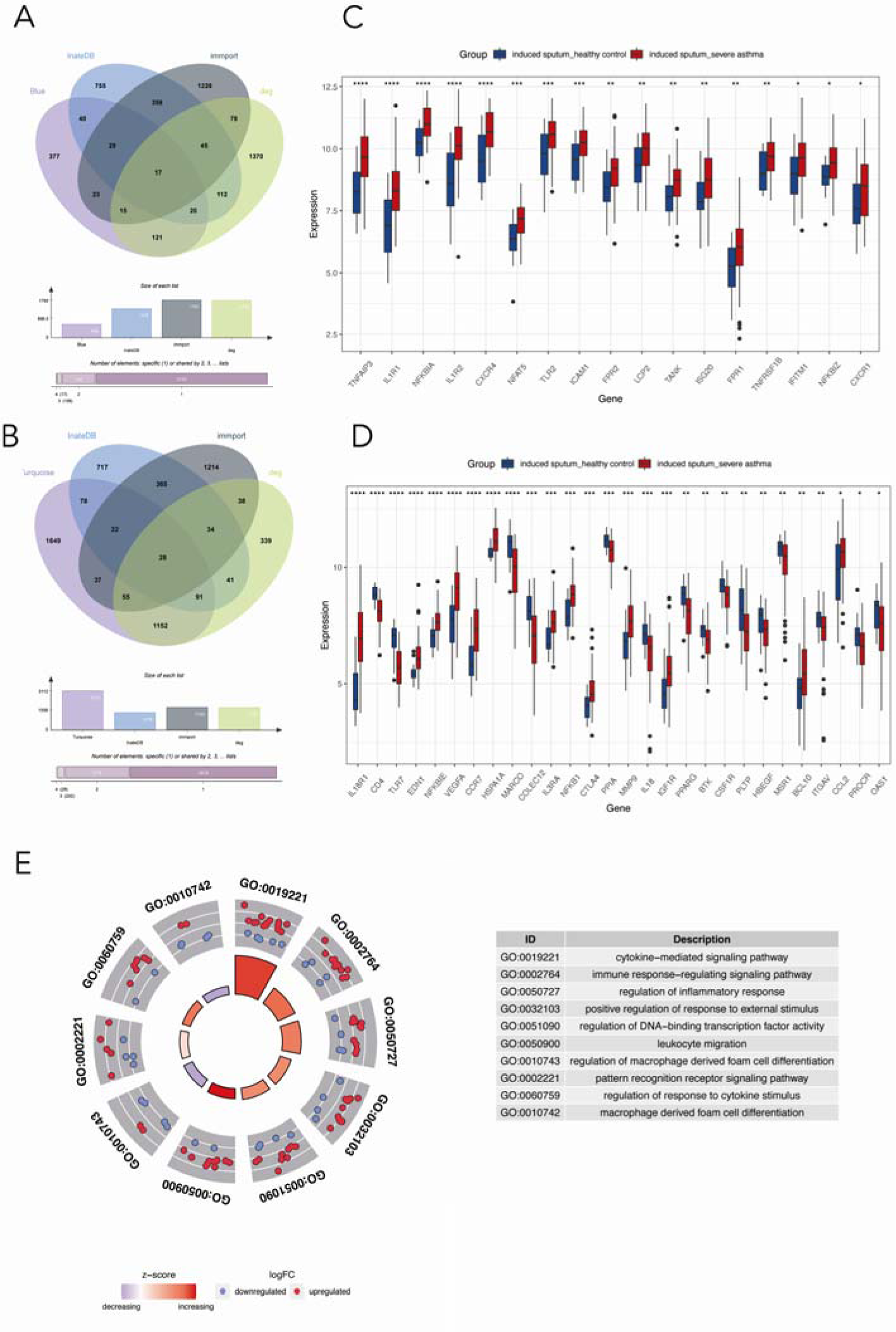
Analysis of immune-related DEGs from the Blue and Turquoise modules and their associations with asthma progression. (A) Venn diagram illustrating the identification of 28 immune-related DEGs shared by the Turquoise module-related genes, DEGs between healthy controls and severe asthma, and immune-related genes from the ImmPort and InnateDB databases. (B) Venn diagram depicting the identification of 17 immune-related DEGs from the Blue module. (C-D) Boxplots displaying the expression levels of the 45 immune-related DEGs in healthy control and severe asthma groups, highlighting the distinct immune features between the two groups. (E) GO enrichment analysis of the 45 immune-related DEGs from the Blue and Turquoise modules, demonstrating significant enrichment in immune and inflammation-related processes, implicating their potential roles in severe asthma pathogenesis.

Our Gene Ontology (GO) enrichment analysis connected these DEGs to immune and inflammatory responses, underscoring their relevance in severe asthma (Figure 4E). The enriched pathways suggest these genes are crucial for immune cell functions, impacting their interactions, regulations, and activations. Our study emphasizes the significant impact of these immune-related DEGs on asthma.

### 3. Development and estimation of machine learning models and Identification and Validation of Key Genes for Severe Asthma Discrimination

To identify the most informative gene features for discriminating between healthy and severe asthma patients, we employed three machine learning algorithms: lasso regression, XGBoost, and random forest models, using 45 immune-related genes from a previous analysis as input features. We randomly divided our sample data into training and testing sets at a 1:1 ratio using caret package and built the three machine learning models on the training set.

We first created a lasso regression model with the glmnet package, selecting features through the lambda.min value (Figure 5A), which simplifies the model by eliminating less critical features. The LASSO model’s efficacy was confirmed on the test set, showcasing an impressive area under the curve (AUC) of 0.94 (Figure 5B). Next, we built an XGBoost model, a choice motivated by its ability to handle sparse data and its robustness against overfitting, using the xgboost package and measured its accuracy with log loss, achieving a 0.93 AUC on the test set (Figure 5C & 5D). Lastly, the random forest model, selected for its versatility and ease of interpretation, highlighted the 30 most important genes, reaching a 0.94 AUC on the test set (Figure 5E & 5F). Higher AUC values indicate superior model performance in distinguishing between the two patient groups.

Integrating the findings from the lasso regression, XGBoost, and random forest algorithms, we identified seven key genes (HBEGF, HSPA1A, CD4, IL18R1, EDN1, CTLA4, IGF1R) that consistently distinguished between healthy and severe asthma cases (Figure 5G). These genes are promising biomarkers for severe asthma, providing insights into the disease’s molecular framework and suggesting potential targets for new treatments. Their relevance to immune responses and inflammation in severe asthma merits further study.

**Figure 5.**
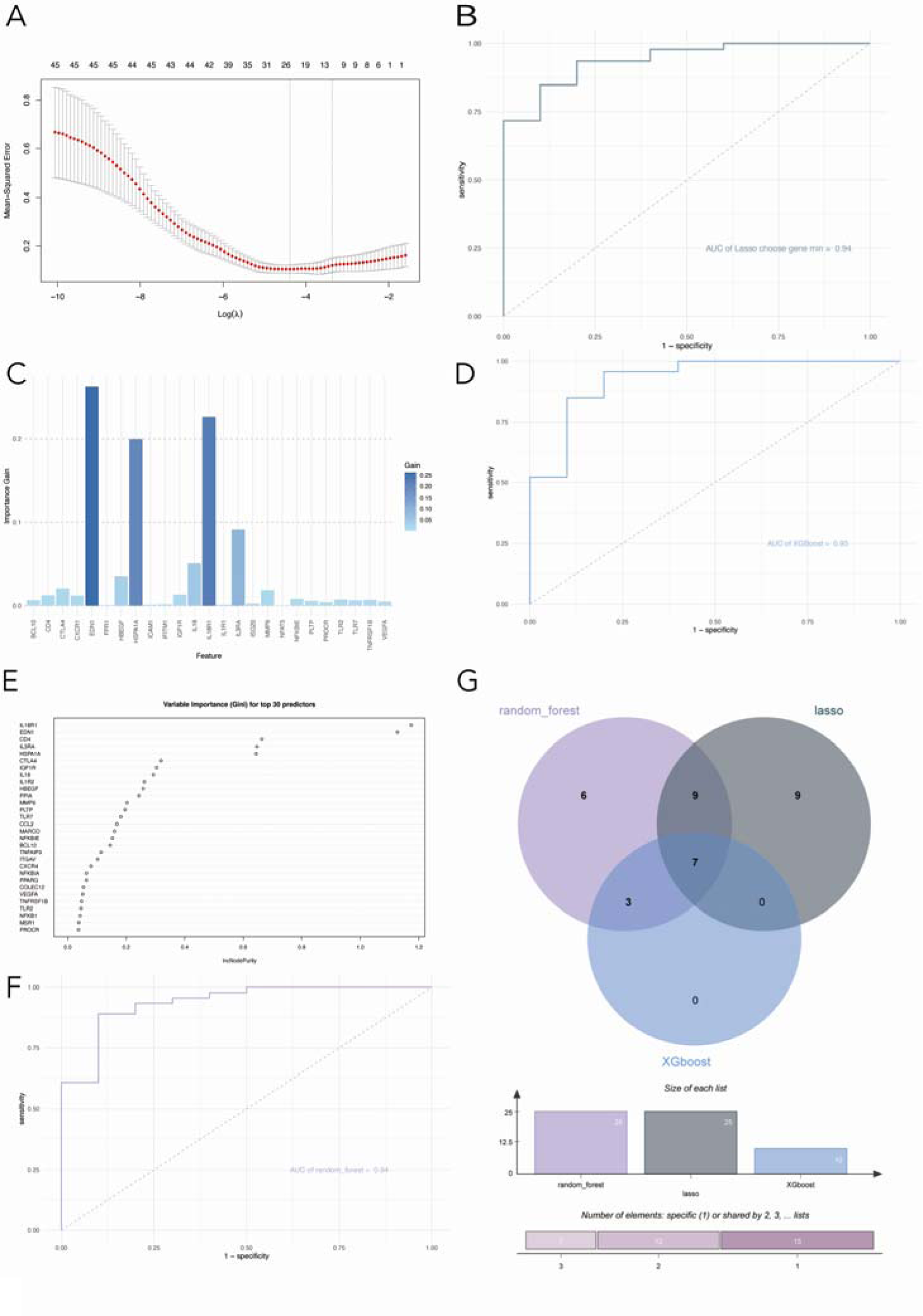
Identification of informative gene features using machine learning algorithms. (A) Lasso regression model construction and feature selection using the glmnet package with lambda.min value. (B) Validation of the Lasso model on the testing set, achieving an AUC of 0.94. (C) XGBoost model construction using the xgboost package with specified parameters and log loss as the evaluation metric. (D) Validation of the XGBoost model on the testing set, achieving an AUC of 0.93. (E) Random forest model construction using the randomForest package and selection of top 30 genes based on importance scores. (F) Validation of the random forest model on the testing set, achieving an AUC of 0.94. Higher AUC values indicate better performance in distinguishing between healthy and severe asthma patients. (G) Identification of key genes for distinguishing between healthy and severe asthma patients. The intersection of three machine learning algorithms—lasso regression, XGBoost, and random forest—reveals seven pivotal genes (HBEGF, HSPA1A, CD4, IL18R1, EDN1, CTLA4, and IGF1R) as robust and discriminative features.

Based on the identified genes, we determined their positions in the human genome (Figure 6A). Subsequently, we utilized receiver operating characteristic (ROC) curves to validate the performance of these seven genes in the test set, validation set, and their combination. The area under the curve (AUC) values demonstrated satisfactory performance across all sets (Figure 6B-H).

**Figure 6.**
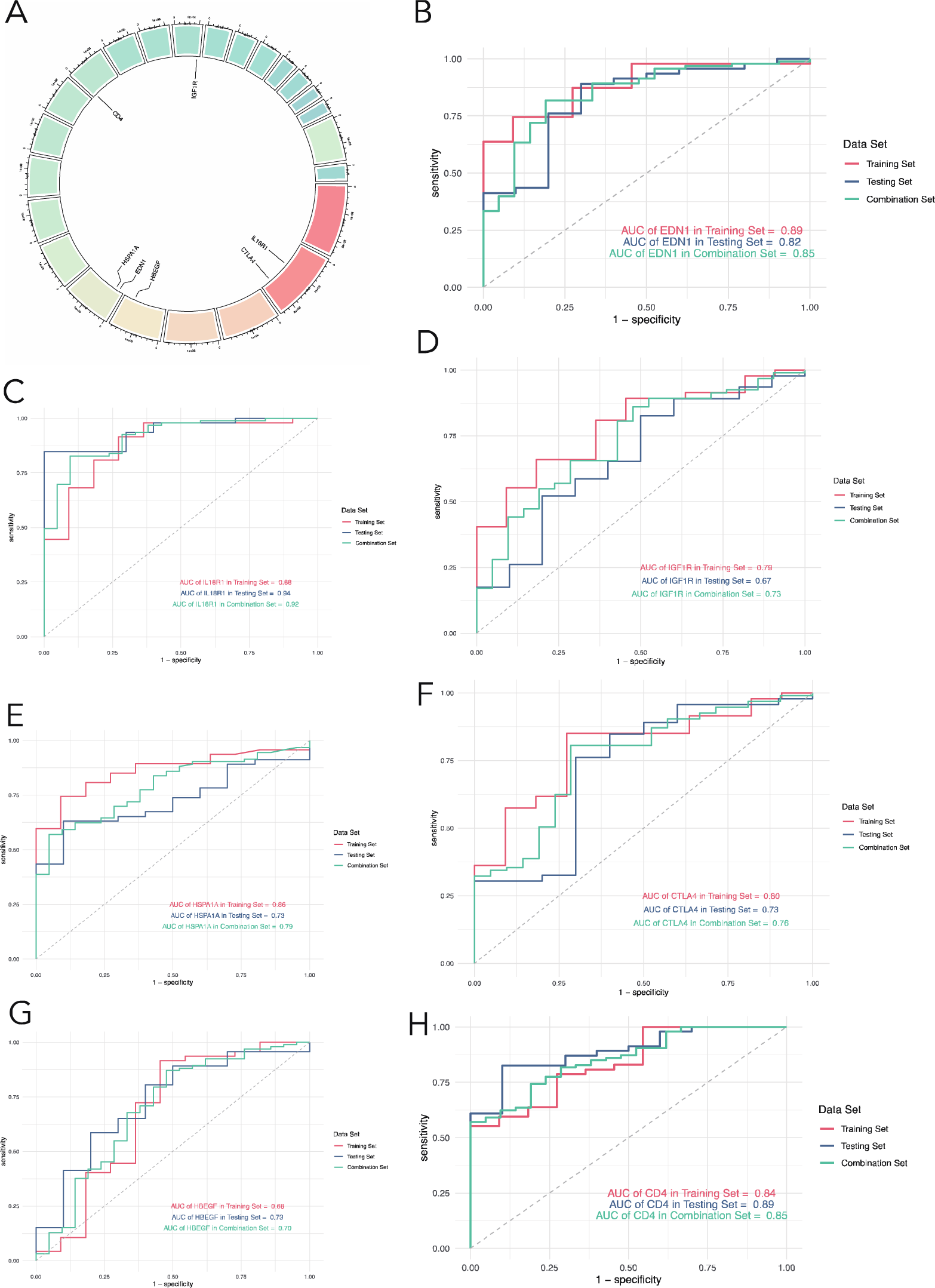
Gene positions and validation performance. (A) Determination of the identified genes’ positions in the human genome. (B-H) Validation of the seven genes’ performance using receiver operating characteristic (ROC) curves in the test set, validation set, and their combination, with area under the curve (AUC) values demonstrating satisfactory performance across all sets.

### 4. Validation of key genes in External Dataset and Animal Model

To corroborate the expression of the seven pivotal genes identified, we delved into transcriptomic samples from healthy controls and severe asthma patients across three external datasets: GSE74986, GSE137268, and GSE74075. However, it’s worth noting that due to microarray probe limitations inherent to the GSE74986 dataset, we couldn’t ascertain the expression levels of all seven genes therein. Regardless, our results from the other datasets and U-BIOPRED consistently demonstrated similar expression trends. Furthermore, the area under the ROC curve validated their effectiveness (Figure 7A-C).

**Figure 7.**
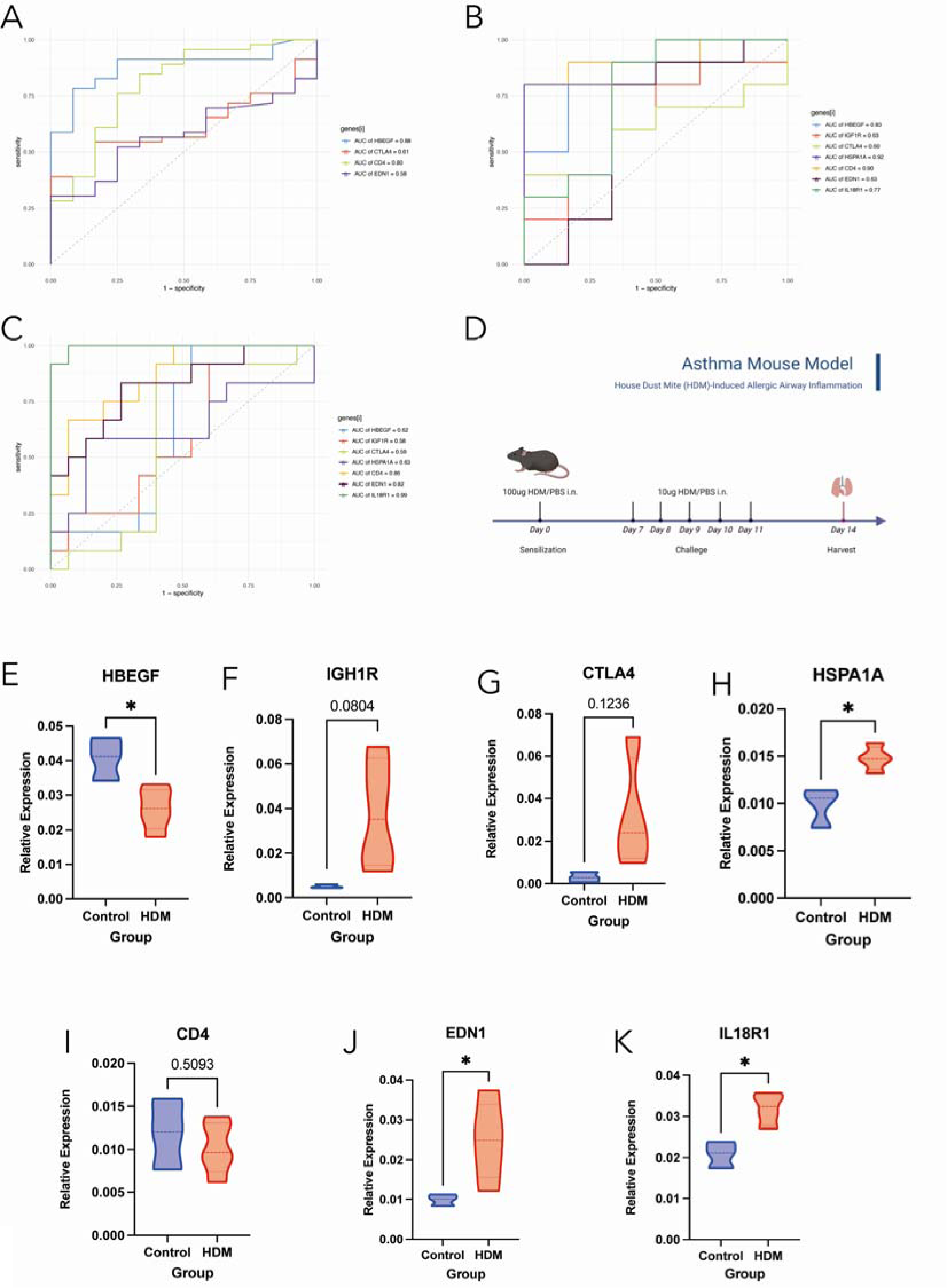
Validation of the seven key genes identified using external datasets and a murine asthma model. **(A-C)** Expression trends and ROC curve analysis of the seven genes in external datasets GSE74986, GSE137268, and GSE74075, demonstrating consistent results with the GSE76262 dataset. (D) Schematic representation of the murine asthma model protocol using house dust mite (HDM) administration. (E-K) Quantitative PCR (qPCR) analysis of the gene expression levels of HBEGF, HSPA1A, CD4, IL18R1, EDN1, CTLA4, and IGF1R in murine lung tissues, showing expression trends consistent with the transcriptomic data in patients and further validating the role of these genes in asthma pathogenesis.

To verify the expression patterns of the seven genes (HBEGF, HSPA1A, CD4, IL18R1, EDN1, CTLA4, and IGF1R) identified by machine learning in asthma pathogenesis, we established a murine model using the following protocol: on day 0, mice were administered 100 µg of house dust mite (HDM) in 40 µL of PBS; on days 7-11, they received 10 µg of HDM in 40 µL of PBS; and on day 14, the mice were sacrificed (Figure 7D).

Subsequently, we isolated RNA from murine lung tissues and performed quantitative PCR (qPCR) analysis to assess gene expression levels. The results demonstrated that the expression trends of HBEGF, HSPA1A, IL18R1, EDN1, CD4, CTLA4, and IGF1R were consistent with those observed at the transcriptomic level in patients (Figure 7E-K and Figure 4).

The consistency between the animal model findings and transcriptomic data supports the reliability of these gene expression patterns as potential biomarkers and therapeutic targets for asthma.

### 5. Identification of immune subtypes of patients with severe asthma

We performed a consensus cluster analysis, in which all severe asthma samples were initially divided into k (k = 2–9) clusters based on the expression matrix of the featured genes above. The cumulative distribution function (CDF) curves of the consensus score matrix and proportion of ambiguous clustering (PAC) statistic indicated that the optimal number was obtained when k = 2 (Figure 8A-C). The same result was achieved from Nbclust testing (Figure 8D).

**Figure 8.**
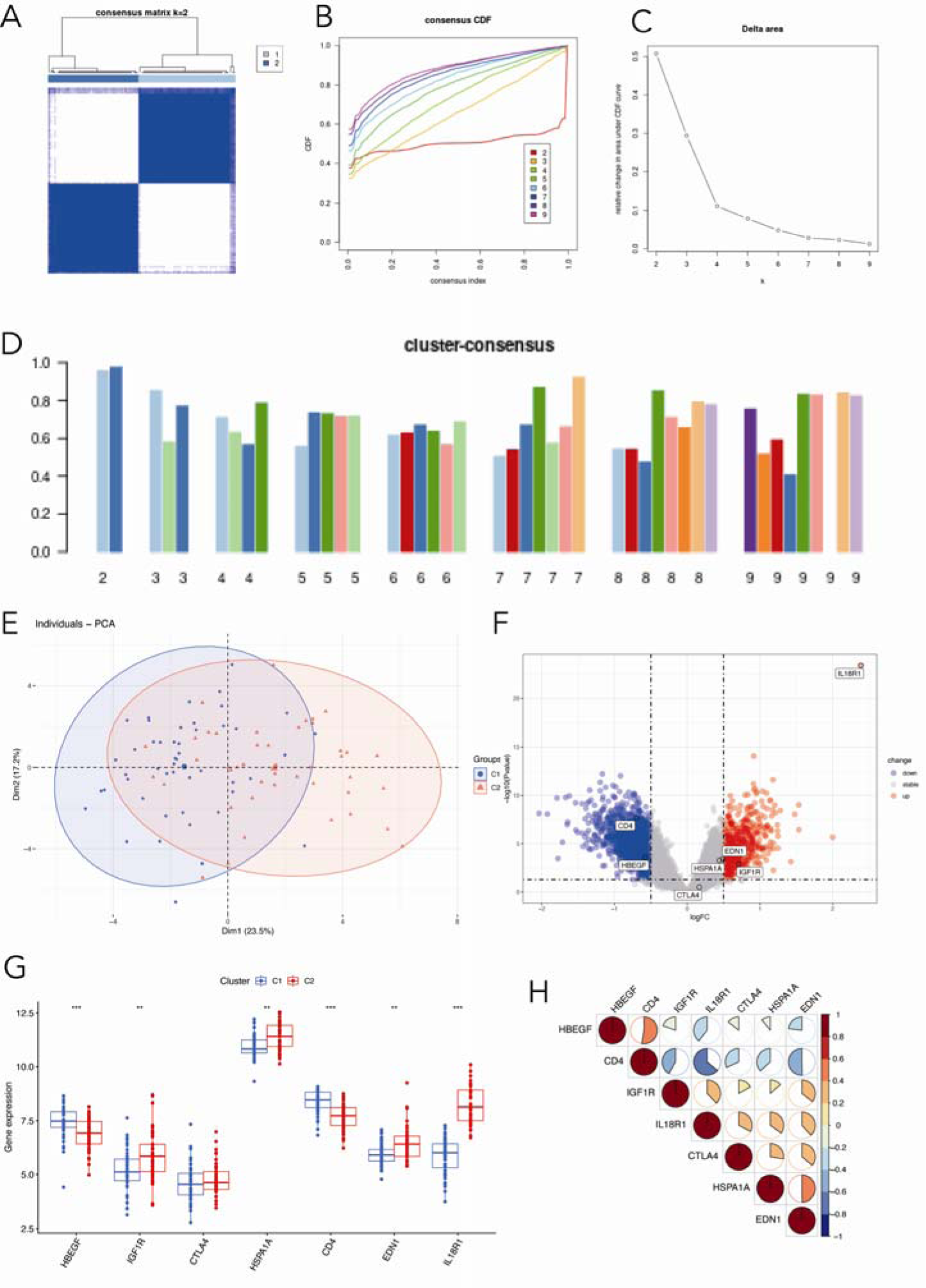
Identification of distinct subtypes within severe asthma patients and their differential gene expression. (A-D) Consensus clustering analysis was performed using the ConsensusClusterPlus package to determine the optimal number of clusters, based on the consensus score matrix, cumulative distribution function (CDF) curve, proportion of ambiguous clustering (PAC) score, and Nbclust. (E) Principal component analysis (PCA) plot illustrating the differences between the two derived subtypes, C1 and C2. (F) Volcano plot showcasing the differentially expressed genes between subtypes C1 and C2. (G) Boxplot depicting the differential expression of the seven critical genes (HBEGF, HSPA1A, CD4, IL18R1, EDN1, CTLA4, and IGF1R) between the C1 and C2 subtypes. (H) Corplot demonstrating the linear relationship between the expression levels of the seven key genes in the subtypes. These results highlight the distinct molecular characteristics present within severe asthma patients, which may have implications for personalized therapeutic approaches.

To visualize the distinctions between the two derived subtypes, we conducted principal component analysis (PCA) using the R function PCA (Figure 8E). The resulting PCA plots effectively displayed the differences between the two subtypes. Subsequently, we executed a differential analysis between subtypes C1 and C2. Although both subtypes consisted of severe asthma patients, a considerable number of differentially expressed genes were observed between them (Figure 8F). This observation was also illustrated in the volcano plot. The seven critical genes exhibited differential expression between the C1 and C2 subtypes, as demonstrated by the boxplot in Figure 8G. Additionally, a corplot revealed a certain degree of linear relationship between the expression levels of these genes (Figure 8H). These findings emphasize the presence of distinct molecular characteristics within severe asthma patients, warranting further exploration to refine personalized therapeutic approaches.

### 6. Differentiation of Immune Characteristics Between Immune-Related Subtypes

To better clarify and understand the biological and immunological differences and relationships between these immune microenvironment subtypes, following the ssGSEA immune infiltration analysis of subtypes C1 and C2, we observed a significant difference in immune cell abundance between the two subgroups, as illustrated in the differential immune cell abundance heatmap (Figure 9A) and the PCA plot (Figure 9B).

**Figure 9.**
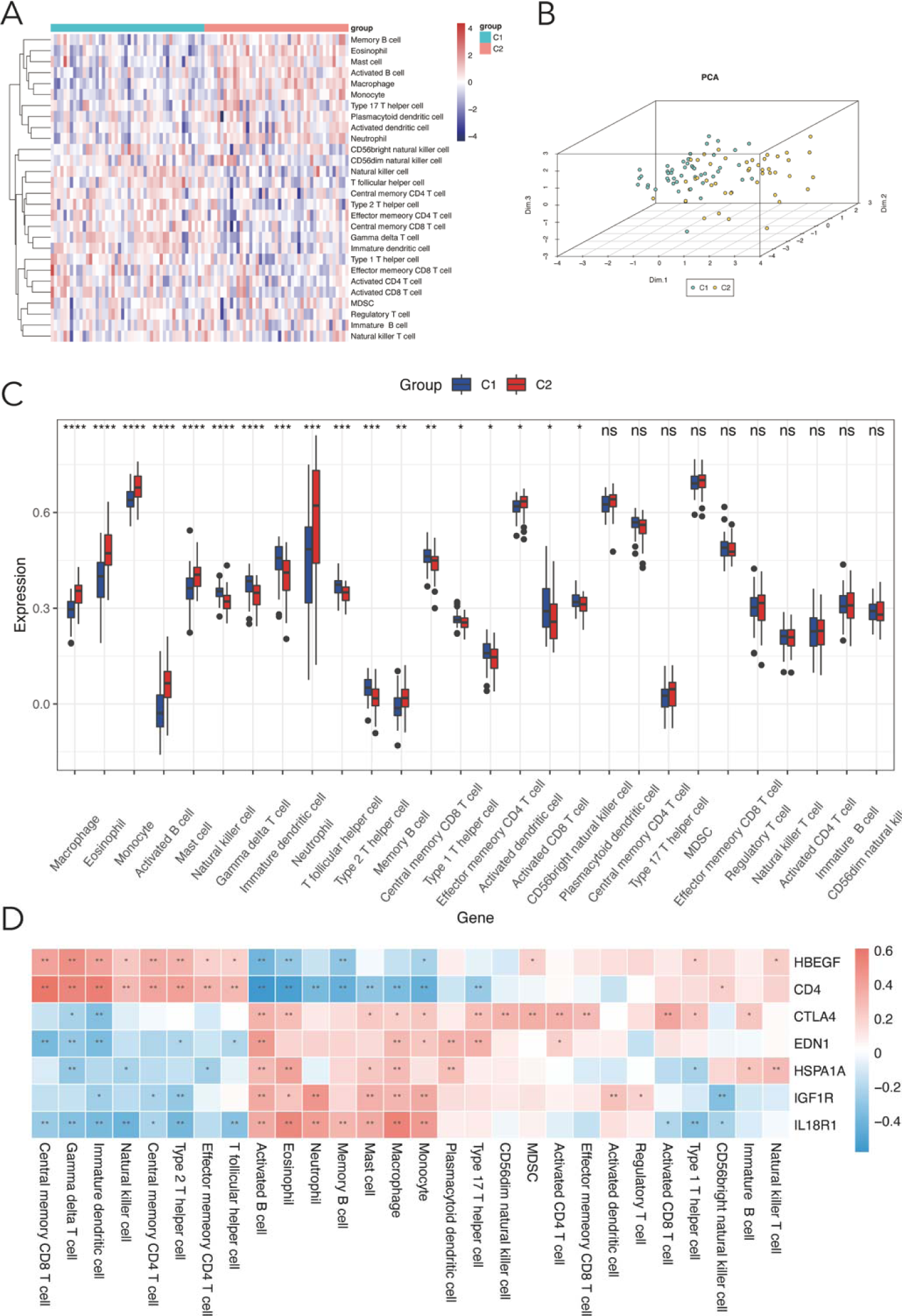
Differential immune cell abundance and correlation with gene expression in subtypes C1 and C2. (A) Heatmap illustrating significant differences in immune cell abundance between subtypes C1 and C2, based on ssGSEA immune infiltration analysis. (B) PCA plot visualizing the distinctions between the two subgroups. (C) Comparison of 28 immune cell subtypes between C1 and C2, with C1 displaying higher infiltration of various T cells, NK cells, and immature DC cells, while C2 is enriched with innate immune cells. (D) Correlation between immune cell abundance from ssGSEA and gene expression, revealing a strong positive correlation between gamma delta T cell abundance and the expression of HBEGF and CD4 in the C1 subtype, and a strong negative correlation with CTLA4, EDN1, HSPA1A, and IL18R1 in the C2 subtype, suggesting the potential role of gamma delta T cells in distinguishing the subtypes and their involvement in differential immune responses.

We further compared the differences in 28 immune cell subtypes between the two subtypes. The C1 subtype exhibited a higher infiltration rate of T cells, including γδ T cells, Tfh cells, Th2 cells, central memory CD8 T cells, effector memory CD4 T cells, and activated CD8 T cells, as well as NK cells and immature DC cells, compared to the C2 subtype. In contrast, the C2 subtype was significantly enriched with innate immune cells such as macrophages, eosinophils, monocytes, activated B cells, mast cells, neutrophils, memory B cells, and activated DC cells(Figure 9C).

These findings imply that the immune response in the C1 subtype is more active, with a stronger adaptive immune response, particularly involving T cells and NK cells. Conversely, the C2 subtype is associated with the activation of innate immune cells. These observations may be linked to distinct asthma pathogenesis mechanisms in the two subtypes. The increased T cell infiltration in the C1 subtype might be associated with Th2 cell activation, a well-established driver of asthma pathogenesis. On the other hand, the activation of innate immune cells in the C2 subtype could be related to environmental factors such as air pollution and allergens. ^34^Further investigation is necessary to elucidate these mechanisms in greater detail.

Subsequently, we explored the correlation between immune cell abundance obtained from ssGSEA and gene expression(Figure 9D). These findings underscore the strong association between the activity of these seven genes and various immune cells, shedding light on the distinct immune landscapes of the two subtypes. This key aspect of our results section highlights the potential of these immune cell signatures in differentiating the C1 and C2 subtypes, as well as the importance of further investigating the role of these genes and immune cells in asthma pathogenesis.

### 7. Function and Pathway Enrichment in Immune-related Subtypes and Therapeutic Target Prediction

To assess disparities in gene expression patterns between the two immune-related subtypes of severe asthma (C1 and C2), we performed a differential expression analysis on gene expression matrices of C1 and C2 compared to healthy controls. We identified 189 upregulated and 306 downregulated differentially expressed genes (DEGs) in the C1 subtype, and 1017 upregulated and 2428 downregulated DEGs in the C2 subtype compared to healthy controls. These findings suggest distinct gene expression profiles in each subtype of severe asthma compared to healthy individuals.

Further analysis of these genes showed that the C1 subtype’s genes are mainly involved in pathways linked to immune responses, such as cytokine interactions, chemokine signaling, and other immune-related activities (Figure 10A, C). In contrast, the genes associated with the C2 subtype are involved in processes related to metabolism, specifically in cellular respiration, mitochondrial function, and protein synthesis (Figure 10B, D). From these findings, we categorized the C1 and C2 subtypes as Immune-Active Subtype (IAS) and Metabolic-Associated Subtype (MAS), respectively. This categorization helps underline the distinct molecular and immune characteristics of each subtype, which could be pivotal in developing customized treatments for severe asthma.

**Figure 10.**
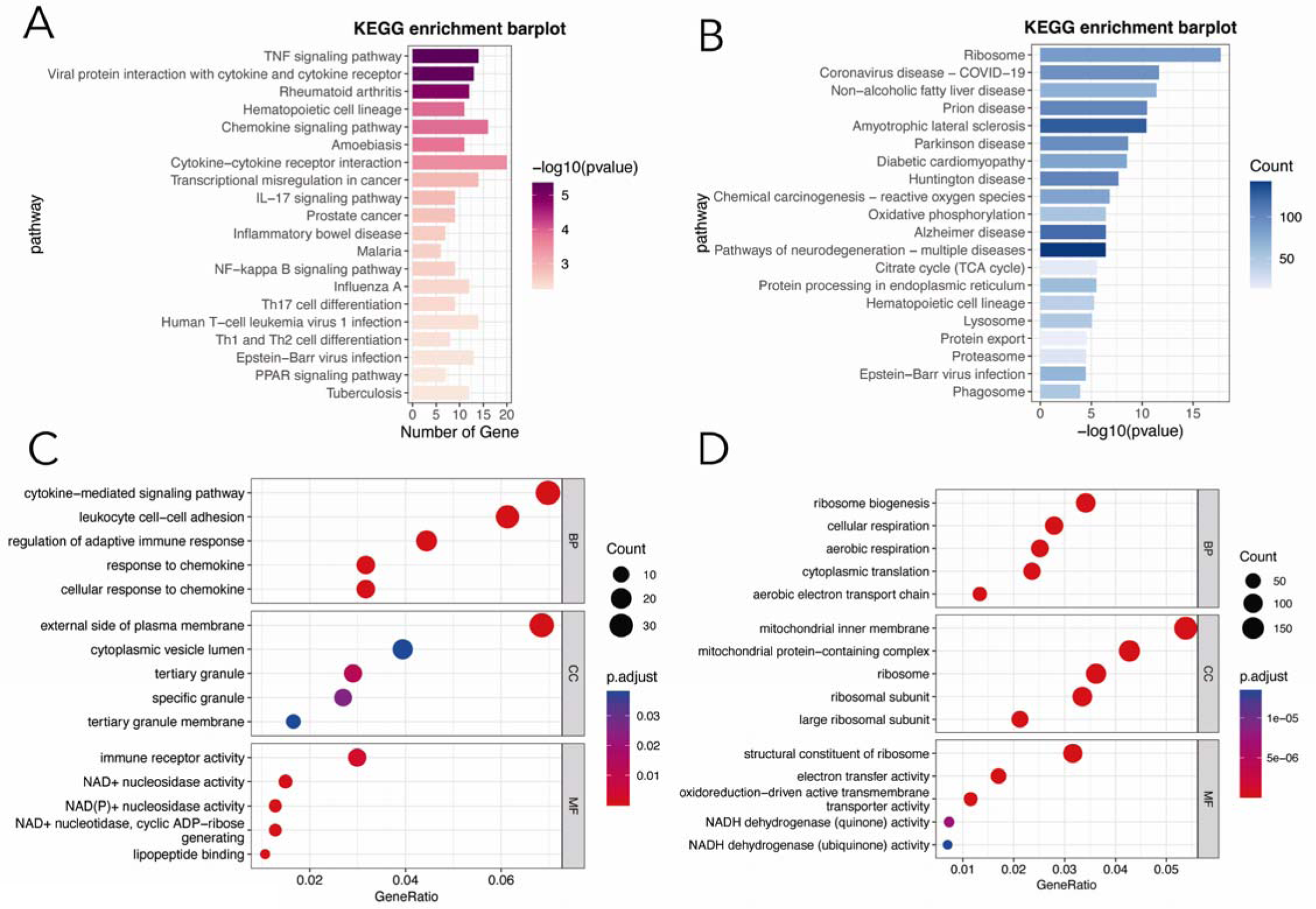
Differential gene expression and enrichment analysis of severe asthma subtypes C1 (IAS) and C2 (MAS). (A, C) Enrichment analysis of C1 subtype DEGs, revealing significant involvement in cytokine-cytokine receptor interaction, chemokine signaling pathway, and GO terms related to cytokine-mediated signaling pathways, leukocyte cell-cell adhesion, and immune receptor activity. (B, D) Enrichment analysis of C2 subtype DEGs, demonstrating a strong association with cellular and aerobic respiration in BP, mitochondrial inner membrane and protein-containing complex in CC, and structural constituents of ribosome and electron transfer activity in MF.

We conducted potential drug target predictions for the top 150 upregulated and downregulated genes in the C1 and C2 subtypes compared to healthy controls using the Cmap database. For the IAS subtype, the top potential targets include FADS3, LAP3, TEAD4, RPS6KB2, and SC5DL (Figure S1A, Supplement Table 1). Meanwhile, the top targets for the MAS subtype are FADS3, ICAM3, HLI-373, TNFRSL1A, and TRA2B (Figure S2B, Supplement Table 2). These suggested targets could guide the development of tailored therapies, offering a step forward in personalized medicine for severe asthma.

## Discussion

Asthma is recognized as a complex disorder with a spectrum of phenotypes beyond the basic allergic and non-allergic categories, requiring detailed study of its various subtypes and the distinct triggers, treatment responses, and pathophysiology associated with each. This nuanced view acknowledges that asthma is not a single condition but rather a constellation of related disorders, each potentially requiring a specialized approach for effective management and therapeutic intervention. Particularly challenging is severe asthma, which involves persistent airway obstruction due to airway-wall remodeling and mucus buildup, highlighting the urgent need for more effective treatments. As we tread further into an era of medical advancement, a myriad of novel treatment methodologies have emerged^19,35^. However, a one-size-fits-all approach is futile. Alarmingly, several treatments, though promising on paper, fall short in efficacy when applied to the real-world heterogeneous patient population. Such observations underline the pressing need for personalized treatment regimens, tailored to the unique genetic and phenotypic profiles of individual patients. Utilizing machine learning and advanced techniques is crucial for developing targeted treatments and gaining a deeper understanding of asthma, especially through examining gene functions and their impact on the disease’s subtypes.

In the evolving field of machine learning for disease research, significant progress has been made. Liu et al. showcased how various analytical methods can create effective predictive models. Xiao, Y. et al. made strides with their use of Convolutional Neural Networks (CNNs) in the Huatuo framework, a breakthrough in understanding gene regulation at the cellular level^36^. Machine learning has notably transformed asthma research. An analysis using the NHANES database found key links between certain factors and pediatric asthma, employing logistic and Lasso regression. This research identified a complex relationship between ethnicity, vitamin B12, sodium, vitamin K, and asthma, providing insights into how nutrition and demographics influence the disease^37^. Furthermore, the discovery of pivotal genes like MMP12, associated with eosinophilic markers, highlights how proteomic analysis can aid asthma diagnosis^38^. However, research hasn’t extensively moved beyond pediatric asthma to include broad datasets and animal model validation. There’s a noticeable absence of studies that merge various machine learning methods to pinpoint asthma-related features. This gap presents an opportunity for groundbreaking research, particularly in personalized medicine, where these methods could significantly tailor interventions to individual needs.

Our study primarily utilized the advanced algorithms of Weighted Gene Co-expression Network Analysis (WGCNA) alongside single-sample Gene Set Enrichment Analysis (ssGSEA) to uncover genes related to immune response in severe asthma. Initial findings showed significant differences in immune cell levels between patients with severe asthma and healthy individuals. By combining WGCNA with machine learning techniques, we developed three models: XGBoost, LASSO, and Random Forest, to efficiently select features and predict severe asthma outcomes. This approach successfully identified seven key genes involved in asthma, validated through various datasets and experimental models. These genes play pivotal roles in the development of asthma by influencing inflammation, airway remodeling, and immune response. HBEGF, part of the epidermal growth factor family, is notable for its increased expression in the epithelium of asthmatic airways, promoting inflammation and sensitivity^39–41^. IGF1R, a gene for a receptor protein in membranes, is involved in asthma development through its elevated expression in asthmatic airway smooth muscle, leading to muscle growth and heightened sensitivity^42,43^. CTLA4, which encodes the cytotoxic T-lymphocyte antigen-4, influences the immune response; its gene variants show diverse links to asthma risk, severity, and treatment outcomes^44^. HSPA1A, a member of the heat shock protein 70 family, contributes to asthma by enhancing airway inflammation via increased production of pro-inflammatory cytokines and chemokines^45,46^. CD4, which encodes a receptor on helper T cells, plays a role in asthma through airway inflammation and sensitivity, with CD4+ T cells releasing pro-inflammatory cytokines when exposed to allergens^47,48^. EDN1, responsible for encoding endothelin-1, a potent vasoconstrictor, is involved in asthma-associated airway inflammation, tightening, and structural changes. Inhibiting its signaling has been shown to reduce sensitivity and inflammation in animal studies^49,50^. IL18R1, coding for the IL-18 cytokine receptor, is associated with inflammation and changes in asthmatic airways^51–54^. Its role is further evidenced by the detection of increased IL18R1 protein levels in lung tissue and the activation of downstream NF-κB and AP-1 pathways, emphasizing the significance of IL-18 signaling in severe asthma^52^. Building on this, our signature, refined by three machine learning methods, demonstrated improved prediction capabilities. Additional validation, based on qRT-PCR results from the HDM asthma animal model, confirmed the unique expression of these genes in asthma’s pathogenesis. This not only reinforced our initial findings but also highlighted the significance of these genes in understanding asthma more broadly.

Leveraging our genomic insights, we identified two distinct immune-related severe asthma subtypes, offering new perspectives on the disease’s complexity. We named these subtypes Immune-Active Subtype (IAS) and Metabolic-Associated Subtype (MAS) for C1 and C2, respectively. IAS features a strong presence of adaptive immune cells, such as various T cell types, highlighting an active inflammatory response typical of asthma^55^. In contrast, MAS is characterized by a significant number of innate immune cells, suggesting a reaction to environmental factors like allergens or pollutants^56^. Our detailed analysis of the immune-related differentially expressed genes (DEGs) further distinguishes these subtypes. IAS is linked with immune cell communication and activation, indicating vigorous immune recruitment. MAS, however, is associated with metabolic functions, hinting at an increased energy demand possibly due to immune responses or changes in airway structure^57–59^. This metabolic emphasis could influence the body’s cellular energy balance and mitochondrial health, both vital in asthma’s development. In summary, IAS is associated with immune response activities, while MAS focuses on metabolic processes. Understanding these differences can guide us in creating more personalized asthma treatments, aiming at the specific pathways of each subtype. However, this is pioneering work, and further research is essential to uncover the detailed roles these pathways play in asthma.

Our groundbreaking findings align with the broader research trends in asthma pathophysiology and classification. The understanding of asthma has significantly evolved, transitioning from traditional categories to more detailed classifications like phenotypes or endotypes^60,61^. This shift highlights the advancement towards precision medicine, where treatment is customized based on detailed evaluations of each patient’s unique characteristics, aiming to enhance care. In asthma classification, a key distinction exists between allergic and eosinophilic asthma^62^. Both are linked to type 2 (T2) inflammation and high eosinophil levels but were traditionally seen as separate^63^. However, treatments targeting T2 inflammation have transformed the management of these asthma types, offering better symptom control for patients with severe cases^14,64,65^. Despite this, current guidelines still recommend different treatments for each type^2,20^, likely due to long-standing views influenced by clinical trial categories. Recognizing the overlap between these classifications is crucial for reevaluating our approach^21,66,67^.

Our study introduces a detailed examination of two immune subtypes in asthma, IAS and MAS, offering fresh insights. We explore the complex nature of asthma by combining immune and metabolic data, contributing new knowledge and suggesting directions for further research. Identifying these subtypes helps refine treatment strategies for severe asthma, enabling personalized care that aligns with each patient’s specific immune patterns. This could also help predict how well patients will respond to certain treatments, with IAS patients potentially benefiting from therapies targeting Th2-driven inflammation, while MAS patients might better respond to treatments focusing on innate immune responses.

The signature of seven immune-related genes offers a promising tool for asthma study, potentially revolutionizing diagnosis and treatment. Easily replicated using a simple PCR-based assay, this signature stands out for its clinical utility. While retrospective analyses have initiated this discovery, future research should focus on prospective, multi-center studies to confirm and expand upon these findings. Despite limitations in the existing public datasets, the identified genes show potential links to critical outcomes. Further investigation into these genes is essential to fully understand their roles and impacts on asthma management and patient care, potentially leading to significant advancements in the field.

In sum, our research provides a refined perspective for understanding and treating asthma, having identified distinct immune subtypes that pave the way for personalized medicine. This breakthrough sets a promising direction for future asthma research, promising to significantly impact ongoing studies and therapeutic approaches. By illuminating the complex nature of asthma, our work underscores the potential for developing interventions that are precisely tailored to individual patient needs, potentially transforming the landscape of asthma care.

## Author Contributions

HY, LY, and PB performed the majority of the analyses, executed the experiments, compiled the data, and contributed equally to this work. TW and HY conceived and orchestrated the study. HY composed the initial manuscript. PB and HY carried out the external validation. HY and XC devised the flowchart. TW scrutinized and refined the manuscript. All authors contributed to the article and endorsed the submitted version.

## Ethics approval and consent to participate

The animal study was reviewed and approved by Ruijin Hospital Animal Ethics Committee.

## Availability of data and materials

The datasets analyzed during the current study are available in the Gene Expression Omnibus (GEO, https://www.ncbi.nlm.nih.gov/geo/). Further inquiries can be directed to the corresponding authors.

## Acknowledgments

The authors would like to thank Desheng Chen and Shuang Tian for their valuable help on the statistical advice.

---

**Supplement Table 1: Top 100 drug target predictions for the Immune-Active Subtype (IAS) in severe asthma patients.**

**Supplement Table 2: Top 100 drug target predictions for the Metabolic-Associated Subtype (MAS) in severe asthma patients.**

**Supplement Table 3: Dataset Overview for Asthma Gene Expression.** This table provides a comprehensive summary of the four datasets used to profile gene expression in asthma research. It includes information on sample sizes, health status of subjects (healthy controls, moderate and severe asthma cases), and the microarray platforms utilized for each dataset, each distinguished by a unique GEO accession number. For further details, such as specific sample counts and the technology behind each microarray platform, refer to Supplement Table S3.

**Supplement Table 4: Primer Sequences for Real-Time RT-PCR Analysis.** This table enumerates the specific primer sequences used for RT-PCR in the study. It details the forward and reverse primer sequences for each gene of interest, including reference genes, to facilitate the quantification of mRNA expression levels in pulmonary tissue.

**Figure S1. Potential Drug Target Recommendations for the IAS and MAS Subtypes.**

