## Supplementary material for "Deciphering the Immune Subtypes and Signature Genes: A Novel Approach Towards Diagnosing and Prognosticating Severe Asthma through Interpretable Machine Learning": Supplenental Table 1

| Rank | Score | Type | ID | Name | Description |
| --- | --- | --- | --- | --- | --- |
| 1 | 99.79 | kd | CGS001-3995 | FADS3 | Fatty acid desaturases |
| 2 | 98.4 | kd | CGS001-51056 | LAP3 | Leucyl aminopeptidase |
| 3 | 97.87 | kd | CGS001-7004 | TEAD4 | - |
| 4 | 97.66 | kd | CGS001-6199 | RPS6KB2 | p70 subfamily |
| 5 | 97.39 | kd | CGS001-6309 | SC5DL | - |
| 6 | 97.11 | cp | BRD-K17349619 | HLI-373 | MDM inhibitor |
| 7 | 96.9 | cp | BRD-A11702965 | chromomycin-a3 | DNA binding agent |
| 8 | 96.77 | kd | CGS001-11014 | KDELR2 | - |
| 9 | 96.76 | kd | CGS001-8233 | ZRSR2 | RNA binding motif (RRM) containing |
| 10 | 96.72 | cp | BRD-K08924299 | palonosetron | Serotonin receptor antagonist |
| 11 | 96.55 | cp | BRD-K89014967 | AS-703026 | MEK inhibitor |
| 12 | 96.34 | kd | CGS001-6812 | STXBP1 | - |
| 13 | 96.25 | cp | BRD-K84085265 | CG-930 | JNK inhibitor |
| 14 | 96.06 | kd | CGS001-10626 | TRIM16 | Tripartite motif containing |
| 15 | 95.91 | oe | ccsbBroad304_06575 | MEF2A | Myocyte enhancer factors |
| 16 | 95.78 | oe | ccsbBroad304_01546 | SLC7A1 | SLC7 family |
| 17 | 95.57 | kd | CGS001-50865 | HEBP1 | Endogenous ligands |
| 18 | 95.53 | kd | CGS001-90589 | ZNF625 | Zinc fingers, C2H2-type |
| 19 | 95.28 | cp | BRD-K39944607 | ochratoxin-a | Phenylalanyl tRNA synthetase inhibitor |
| 20 | 95.19 | oe | ccsbBroad304_07699 | GPR83 | GPCR / Class A : Orphans |
| 21 | 95.16 | kd | CGS001-23196 | FAM120A | - |
| 22 | 95.11 | kd | CGS001-54107 | POLE3 | DNA polymerases |
| 23 | 95.04 | kd | CGS001-8508 | NIPSNAP1 | - |
| 24 | 95.04 | kd | CGS001-3456 | IFNB1 | Interferons |
| 25 | 94.91 | kd | CGS001-8864 | PER2 | - |
| 26 | 94.69 | kd | CGS001-91137 | SLC25A46 | Miscellaneous SLC25 mitochondrial transporters |
| 27 | 94.51 | cp | BRD-K49456190 | prima-1-met | thioredoxin inhibitor |
| 28 | 94.51 | kd | CGS001-55423 | SIRPG | Immunoglobulin superfamily / C1-set domain containing |
| 29 | 94.22 | cp | BRD-K68191783 | ALW-II-38-3 | Ephrin inhibitor |
| 30 | 94.1 | kd | CGS001-486 | FXYD2 | Ion transport regulator |
| 31 | 93.95 | cc |  | Akt Signaling GOF | - |
| 32 | 93.91 | kd | CGS001-8835 | SOCS2 | SH2 domain containing |
| 33 | 93.71 | kd | CGS001-8717 | TRADD | - |
| 34 | 93.7 | kd | CGS001-10269 | ZMPSTE24 | - |
| 35 | 93.68 | cp | BRD-M16762496 | PIK-75 | DNA protein kinase inhibitor |
| 36 | 93.53 | kd | CGS001-894 | CCND2 | - |
| 37 | 93.52 | oe | ccsbBroad304_08461 | CDCA4 | - |
| 38 | 93.5 | kd | CGS001-23077 | MYCBP2 | - |
| 39 | 93.5 | kd | CGS001-55869 | HDAC8 | Histone deacetylases |
| 40 | 93.41 | oe | ccsbBroad304_06089 | DDIT3 | - |
| 41 | 93.16 | kd | CGS001-134187 | POU5F2 | Homeoboxes / POU class |
| 42 | 93.11 | kd | CGS001-1387 | CREBBP | Chromatin-modifying enzymes / K-acetyltransferases |
| 43 | 93.07 | kd | CGS001-26136 | TES | - |
| 44 | 93.04 | kd | CGS001-6342 | SCP2 | - |
| 45 | 93 | kd | CGS001-6434 | TRA2B | RNA binding motif (RRM) containing |
| 46 | 92.95 | kd | CGS001-29775 | CARD10 | - |
| 47 | 92.63 | kd | CGS001-22908 | SACM1L | - |
| 48 | 92.36 | kd | CGS001-648 | BMI1 | Polycomb group ring fingers |
| 49 | 92.27 | oe | ccsbBroad304_06338 | GTF2A2 | General transcription factors |
| 50 | 92.18 | kd | CGS001-1390 | CREM | basic leucine zipper proteins |
| 51 | 92.16 | kd | CGS001-79733 | E2F8 | - |
| 52 | 92.09 | oe | ccsbBroad304_07006 | ST14 | Serine peptidases / Transmembrane |
| 53 | 91.8 | oe | ccsbBroad304_00046 | AKT1 | Akt (Protein kinase B) |
| 54 | 91.78 | kd | CGS001-6541 | SLC7A1 | SLC7 family |
| 55 | 91.59 | kd | CGS001-2168 | FABP1 | Fatty acid-binding proteins |
| 56 | 91.2 | oe | ccsbBroad304_06847 | RBBP4 | WD repeat domain containing |
| 57 | 91.01 | kd | CGS001-25805 | BAMBI | - |
| 58 | 90.92 | kd | CGS001-114112 | TXNRD3 | - |
| 59 | 90.89 | kd | CGS001-11113 | CIT | Other DMPK family kinases |
| 60 | 90.5 | kd | CGS001-10733 | PLK4 | Polo-like kinase (PLK) family |
| 61 | 90.46 | cp | BRD-K50720187 | flupirtine | Glutamate receptor antagonist |
| 62 | 90.42 | cp | BRD-K37720887 | SB-525334 | TGF beta receptor inhibitor |
| 63 | 90.21 | kd | CGS001-1385 | CREB1 | basic leucine zipper proteins |
| 64 | 90 | oe | ccsbBroad304_00294 | CETN3 | EF-hand domain containing |
| 65 | 89.81 | kd | CGS001-6660 | SOX5 | SRY (sex determining region Y)-boxes |
| 66 | 89.57 | kd | CGS001-6925 | TCF4 | Basic helix-loop-helix proteins |
| 67 | 89.48 | kd | CGS001-134864 | TAAR1 | GPCR / Class A : Trace amine associated receptors |
| 68 | 89.39 | kd | CGS001-10652 | YKT6 | - |
| 69 | 89.2 | kd | CGS001-2288 | FKBP4 | Tetratricopeptide (TTC) repeat domain containing |
| 70 | 89.18 | cp | BRD-A13122391 | triptolide | RNA polymerase inhibitor |
| 71 | 88.96 | kd | CGS001-54805 | CNNM2 | - |
| 72 | 88.88 | kd | CGS001-79145 | CHCHD7 | Coiled-coil-helix-coiled-coil-helix domain containing |
| 73 | 88.77 | kd | CGS001-4125 | MAN2B1 | - |
| 74 | 88.63 | oe | ccsbBroad304_06130 | DUSP4 | Protein tyrosine phosphatases / Class I Cys-based PTPs : MAP kinase phosphatases |
| 75 | 88.6 | kd | CGS001-85377 | MICALL1 | - |
| 76 | 88.34 | kd | CGS001-10957 | PNRC1 | - |
| 77 | 88.32 | cp | BRD-A14985772 | ascorbyl-palmitate | antioxidant |
| 78 | 88.31 | kd | CGS001-4204 | MECP2 | - |
| 79 | 88.28 | kd | CGS001-4712 | NDUFB6 | Mitochondrial respiratory chain complex / Complex I |
| 80 | 87.94 | cp | BRD-K04146668 | GW-441756 | Growth factor receptor inhibitor |
| 81 | 87.93 | kd | CGS001-9925 | ZBTB5 | BTB/POZ domain containing |
| 82 | 87.86 | cp | BRD-K05804044 | AZ-628 | RAF inhibitor |
| 83 | 87.81 | kd | CGS001-7159 | TP53BP2 | Ankyrin repeat domain containing |
| 84 | 87.72 | cp | BRD-K05104363 | PD-184352 | MEK inhibitor |
| 85 | 87.57 | cp | BRD-K01638814 | rilmenidine | Adrenergic receptor agonist |
| 86 | 87.52 | cp | BRD-K64785675 | TG100-115 | -666 |
| 87 | 87.5 | oe | ccsbBroad304_02708 | TRIM32 | - |
| 88 | 87.31 | kd | CGS001-51128 | SAR1B | - |
| 89 | 87.15 | kd | CGS001-51335 | NGRN | - |
| 90 | 86.94 | kd | CGS001-60468 | BACH2 | BTB/POZ domain containing |
| 91 | 86.87 | kd | CGS001-11319 | ECD | - |
| 92 | 86.85 | kd | CGS001-6793 | STK10 | SLK subfamily |
| 93 | 86.82 | kd | CGS001-27336 | HTATSF1 | RNA binding motif (RRM) containing |
| 94 | 86.77 | kd | CGS001-83440 | ADPGK | - |
| 95 | 86.49 | cp | BRD-K01555864 | dibenzoylmethane | Antineoplastic |
| 96 | 86.44 | kd | CGS001-994 | CDC25B | Protein tyrosine phosphatases / Class III Cys-based PTPs |
| 97 | 86.4 | kd | CGS001-1001 | CDH3 | Cadherins / Major cadherins |
| 98 | 86.2 | kd | CGS001-6519 | SLC3A1 | SLC3 family |
| 99 | 86.07 | oe | ccsbBroad304_02771 | HEY1 | Basic helix-loop-helix proteins |
| 100 | 85.88 | cp | BRD-K67013324 | luzindole | Melatonin receptor antagonist |
