## Supplementary material for "Deciphering the Immune Subtypes and Signature Genes: A Novel Approach Towards Diagnosing and Prognosticating Severe Asthma through Interpretable Machine Learning": Supplenental Table 2

| Rank | Score | Type | ID | Name | Description |
| --- | --- | --- | --- | --- | --- |
| 1 | 99.89 | kd | CGS001-3995 | FADS3 | Fatty acid desaturases |
| 2 | 99.58 | kd | CGS001-3385 | ICAM3 | CD molecules |
| 3 | 99.51 | cp | BRD-K17349619 | HLI-373 | MDM inhibitor |
| 4 | 99.42 | kd | CGS001-7132 | TNFRSF1A | Tumour necrosis factor (TNF) receptor family |
| 5 | 99.17 | kd | CGS001-6434 | TRA2B | RNA binding motif (RRM) containing |
| 6 | 98.96 | kd | CGS001-6199 | RPS6KB2 | p70 subfamily |
| 7 | 98.95 | kd | CGS001-54107 | POLE3 | DNA polymerases |
| 8 | 98.74 | kd | CGS001-23039 | XPO7 | Exportins |
| 9 | 98.72 | kd | CGS001-10269 | ZMPSTE24 | - |
| 10 | 98.55 | kd | CGS001-4352 | MPL | Prolactin receptor family |
| 11 | 98.31 | cp | BRD-K84085265 | CG-930 | JNK inhibitor |
| 12 | 98.31 | cp | BRD-K74236984 | UNC-0321 | Histone lysine methyltransferase inhibitor |
| 13 | 98.28 | kd | CGS001-25805 | BAMBI | - |
| 14 | 98.13 | kd | CGS001-90589 | ZNF625 | Zinc fingers, C2H2-type |
| 15 | 98.07 | kd | CGS001-54805 | CNNM2 | - |
| 16 | 98.05 | kd | CGS001-5734 | PTGER4 | GPCR / Class A : Prostanoid receptors |
| 17 | 97.97 | kd | CGS001-648 | BMI1 | Polycomb group ring fingers |
| 18 | 97.72 | kd | CGS001-23013 | SPEN | RNA binding motif (RRM) containing |
| 19 | 97.48 | kd | CGS001-8312 | AXIN1 | Serine/threonine phosphatases / Protein phosphatase 1, regulatory subunits |
| 20 | 97.47 | oe | ccsbBroad304_02771 | HEY1 | Basic helix-loop-helix proteins |
| 21 | 97.27 | cp | BRD-A13122391 | triptolide | RNA polymerase inhibitor |
| 22 | 97.21 | kd | CGS001-8835 | SOCS2 | SH2 domain containing |
| 23 | 97.01 | kd | CGS001-10957 | PNRC1 | - |
| 24 | 96.83 | cp | BRD-K39944607 | ochratoxin-a | Phenylalanyl tRNA synthetase inhibitor |
| 25 | 96.76 | kd | CGS001-9128 | PRPF4 | WD repeat domain containing |
| 26 | 96.6 | cp | BRD-K32330832 | VER-155008 | HSP inhibitor |
| 27 | 96.47 | kd | CGS001-7424 | VEGFC | - |
| 28 | 96.46 | kd | CGS001-10019 | SH2B3 | Pleckstrin homology (PH) domain containing |
| 29 | 96.44 | cp | BRD-K30677119 | PP-30 | RAF inhibitor |
| 30 | 96.33 | kd | CGS001-9189 | ZBED1 | Pseudoautosomal regions / PAR1 |
| 31 | 96.3 | oe | ccsbBroad304_07699 | GPR83 | GPCR / Class A : Orphans |
| 32 | 96.29 | cp | BRD-K41895714 | AS-605240 | PI3K inhibitor |
| 33 | 96.25 | cp | BRD-K89014967 | AS-703026 | MEK inhibitor |
| 34 | 96.18 | kd | CGS001-6342 | SCP2 | - |
| 35 | 96.14 | kd | CGS001-79143 | MBOAT7 | - |
| 36 | 96.02 | kd | CGS001-5930 | RBBP6 | - |
| 37 | 95.84 | kd | CGS001-25874 | BRP44 | - |
| 38 | 95.81 | oe | ccsbBroad304_06575 | MEF2A | Myocyte enhancer factors |
| 39 | 95.61 | kd | CGS001-10308 | ZNF267 | Zinc fingers, C2H2-type |
| 40 | 95.6 | oe | ccsbBroad304_01953 | PIAS1 | Zinc fingers, MIZ-type |
| 41 | 95.56 | oe | ccsbBroad304_07245 | PPFIBP2 | Sterile alpha motif (SAM) domain containing |
| 42 | 95.32 | oe | ccsbBroad304_07294 | TNFSF10 | CD molecules |
| 43 | 95.14 | oe | ccsbBroad304_02738 | RRS1 | - |
| 44 | 95.11 | oe | ccsbBroad304_13952 | SHMT2 | - |
| 45 | 95.01 | oe | ccsbBroad304_06483 | KCNK1 | Two-P potassium channels |
| 46 | 94.95 | kd | CGS001-3459 | IFNGR1 | Interferon receptor family |
| 47 | 94.93 | kd | CGS001-7004 | TEAD4 | - |
| 48 | 94.87 | kd | CGS001-7159 | TP53BP2 | Ankyrin repeat domain containing |
| 49 | 94.83 | kd | CGS001-1810 | DR1 | - |
| 50 | 94.65 | kd | CGS001-163486 | DENND1B | DENN/MADD domain containing |
| 51 | 94.58 | cp | BRD-A11702965 | chromomycin-a3 | DNA binding agent |
| 52 | 94.52 | cp | BRD-K08924299 | palonosetron | Serotonin receptor antagonist |
| 53 | 94.42 | kd | CGS001-6925 | TCF4 | Basic helix-loop-helix proteins |
| 54 | 94.3 | cp | BRD-K50720187 | flupirtine | Glutamate receptor antagonist |
| 55 | 94.14 | cp | BRD-K09668667 | benzo(a)pyrene | Carcinogen |
| 56 | 94.13 | kd | CGS001-26136 | TES | - |
| 57 | 93.93 | kd | CGS001-9261 | MAPKAPK2 | MAPKAPK subfamily |
| 58 | 93.73 | kd | CGS001-23621 | BACE1 | Pepsin |
| 59 | 93.66 | kd | CGS001-6883 | TAF12 | - |
| 60 | 93.37 | kd | CGS001-344 | APOC2 | Apolipoproteins |
| 61 | 93.34 | kd | CGS001-2168 | FABP1 | Fatty acid-binding proteins |
| 62 | 93.11 | kd | CGS001-3586 | IL10 | Interleukins and interleukin receptors |
| 63 | 93.1 | oe | ccsbBroad304_06338 | GTF2A2 | General transcription factors |
| 64 | 92.99 | oe | ccsbBroad304_00047 | AKT2 | Akt (Protein kinase B) |
| 65 | 92.73 | oe | ccsbBroad304_03451 | SLC35F2 | SLC35 family of nucleotide sugar transporters |
| 66 | 92.58 | oe | ccsbBroad304_02845 | PRPF6 | - |
| 67 | 92.57 | kd | CGS001-466 | ATF1 | basic leucine zipper proteins |
| 68 | 92.39 | cp | BRD-U44618005 | WH-4023 | SRC inhibitor |
| 69 | 92.38 | cp | BRD-U73238814 | QL-XI-92 | DDR1 inhibitor |
| 70 | 92.35 | kd | CGS001-9925 | ZBTB5 | BTB/POZ domain containing |
| 71 | 92.2 | kd | CGS001-56940 | DUSP22 | Protein tyrosine phosphatases / Class I Cys-based PTPs : Atypical dual specificity phosphatases |
| 72 | 92.13 | kd | CGS001-11124 | FAF1 | UBX domain containing |
| 73 | 92.06 | kd | CGS001-25925 | ZNF521 | Zinc fingers, C2H2-type |
| 74 | 92 | kd | CGS001-5970 | RELA | NFkappaB transcription factor family |
| 75 | 91.81 | kd | CGS001-3035 | HARS | Aminoacyl tRNA synthetases / Class II |
| 76 | 91.78 | kd | CGS001-4712 | NDUFB6 | Mitochondrial respiratory chain complex / Complex I |
| 77 | 91.72 | kd | CGS001-26292 | MYCBP | - |
| 78 | 91.66 | kd | CGS001-10652 | YKT6 | - |
| 79 | 91.56 | kd | CGS001-1381 | CRABP1 | Fatty acid-binding proteins |
| 80 | 91.53 | cp | BRD-K64451768 | GANT-58 | GLI antagonist |
| 81 | 91.43 | kd | CGS001-6397 | SEC14L1 | - |
| 82 | 91.22 | oe | ccsbBroad304_06847 | RBBP4 | WD repeat domain containing |
| 83 | 91.14 | kd | CGS001-22908 | SACM1L | - |
| 84 | 91.07 | kd | CGS001-134864 | TAAR1 | GPCR / Class A : Trace amine associated receptors |
| 85 | 90.96 | kd | CGS001-688 | KLF5 | Kruppel-like transcription factors |
| 86 | 90.83 | kd | CGS001-5771 | PTPN2 | Protein tyrosine phosphatases |
| 87 | 90.82 | kd | CGS001-80347 | COASY | - |
| 88 | 90.8 | kd | CGS001-7181 | NR2C1 | Testicular receptors |
| 89 | 90.75 | kd | CGS001-60468 | BACH2 | BTB/POZ domain containing |
| 90 | 90.61 | kd | CGS001-6812 | STXBP1 | - |
| 91 | 90.02 | kd | CGS001-79733 | E2F8 | - |
| 92 | 90 | kd | CGS001-29916 | SNX11 | Sorting nexins |
| 93 | 89.8 | kd | CGS001-3597 | IL13RA1 | IL-2 receptor family |
| 94 | 89.79 | kd | CGS001-27336 | HTATSF1 | RNA binding motif (RRM) containing |
| 95 | 89.77 | kd | CGS001-23325 | KIAA1033 | - |
| 96 | 89.68 | cp | BRD-K33483813 | actarit | Interleukin receptor agonist |
| 97 | 89.34 | cp | BRD-K01192156 | tyrphostin-AG-112 | Protein tyrosine kinase inhibitor |
| 98 | 89.14 | oe | ccsbBroad304_07006 | ST14 | Serine peptidases / Transmembrane |
| 99 | 88.87 | oe | ccsbBroad304_02276 | MVP | - |
| 100 | 88.77 | kd | CGS001-5442 | POLRMT | - |
