## Supplementary material for "Deciphering the Immune Subtypes and Signature Genes: A Novel Approach Towards Diagnosing and Prognosticating Severe Asthma through Interpretable Machine Learning": Supplenental Table 4

|  | **Primers Implemented for RT-PCR Analysis** | |  |
| --- | --- | --- | --- |
|  | **GENE** | **Forward** | **Reverse** |
|  | HBEGF | CGGGGAGTGCAGATACCTG | TTCTCCACTGGTAGAGTCAGC |
|  | GAPDH | AGGTCGGTGTGAACGGATTTG | GGGGTCGTTGATGGCAACA |
|  | IGF1R | GTGGGGGCTCGTGTTTCTC | GATCACCGTGCAGTTTTCCA |
|  | CTLA4 | TTTTGTAGCCCTGCTCACTCT | CTGAAGGTTGGGTCACCTGTA |
|  | HSPA1A | TGGTGCAGTCCGACATGAAG | GCTGAGAGTCGTTGAAGTAGGC |
|  | CD4 | TCCTAGCTGTCACTCAAGGGA | TCAGAGAACTTCCAGGTGAAGA |
|  | EDN1 | GCACCGGAGCTGAGAATGG | GTGGCAGAAGTAGACACACTC |
|  | IL18R1 | TCACCGATCACAAATTCATGTGG | TGGTGGCTGTTTCATTCCTGT |
