## Supplementary material for "Deciphering the Immune Subtypes and Signature Genes: A Novel Approach Towards Diagnosing and Prognosticating Severe Asthma through Interpretable Machine Learning": Supplenental Table 3

| **Comparative Overview of Asthma Gene Expression Datasets** | | |  |  |  |  |
| --- | --- | --- | --- | --- | --- | --- |
| Dataset | GEO Accession | Sample Source | Healthy Controls | Moderate Asthma Cases | Severe Asthma Cases | Platform |
| U-BIOPRED | GSE76262 | Induced Sputum | 21 | 25 | 93 | Affymetrix U133 Plus 2.0 PM-only arrays |
| Arron JR et al. | GSE74986 | Not Specified | 12 | - | 46 | Agilent-014850 Whole Human Genome Microarray |
| Li Q et al. | GSE74075 | Sputum | 6 | - | 10 | Illumina Sentrix HumanRef-8 V3 Expression BeadChips |
| Baines KJ et al. | GSE137268 | Induced Sputum | 15 | - | 54 | Illumina HumanRef-8 v2.0 Expression BeadChip |
